# Variational Inference Using Approximate Likelihood Under the Coalescent With Recombination

**DOI:** 10.1101/2020.08.19.258137

**Authors:** Xinhao Liu, Huw A. Ogilvie, Luay Nakhleh

## Abstract

Coalescent methods are proven and powerful tools for population genetics, phylogenetics, epidemiology, and other fields. A promising avenue for the analysis of large genomic alignments, which are increasingly common, are coalescent hidden Markov model (coalHMM) methods, but these methods have lacked general usability and flexibility. We introduce a novel method for automatically learning a coalHMM and inferring the posterior distributions of evolutionary parameters using black-box variational inference, with the transition rates between local genealogies derived empirically by simulation. This derivation enables our method to work directly with three or four taxa and through a divide-and-conquer approach with more taxa. Using a simulated data set resembling a human-chimp-gorilla scenario, we show that our method has comparable or better accuracy to previous coalHMM methods. Both species divergence times and population sizes were accurately inferred. The method also infers local genealogies and we report on their accuracy. Furthermore, we illustrate how to scale the method to larger data sets through a divide-and-conquer approach. This accuracy means our method is useful now, and by deriving transition rates by simulation it is flexible enough to enable future implementations of all kinds of population models.

## Introduction

A powerful and widely accepted and employed mathematical framework for capturing the evolution of genomes and their individual loci is the *theory of coalescence* (Kingman, 1982). This framework, applied to the increasingly available genomic data, has “turned theoretical population genetics on its head” (Hartl and Clark, 2007) and propelled population and phylogenomic inferences to successful applications that span several fields of biology and biomedicine (Siepel, 2009; Rogers and Gibbs, 2014). Coalescent-based models allow for estimating the values of parameters, including population divergence times, mutation and recombination rates, ancestral population sizes, population structure, etc., from patterns of site frequencies and local genealogies (Hartl and Clark, 2007; Nielsen and Slatkin, 2013).

To account for varying levels of complexities in evolutionary histories, the standard coalescent has been extended in various directions to accommodate processes such as recombination, population structure and migration, and selection (Hudson, 1990; Wakeley, 2008). In this work, we focus on the coalescent with recombination, illustrated in Fig. 1a.

**Figure 1:**
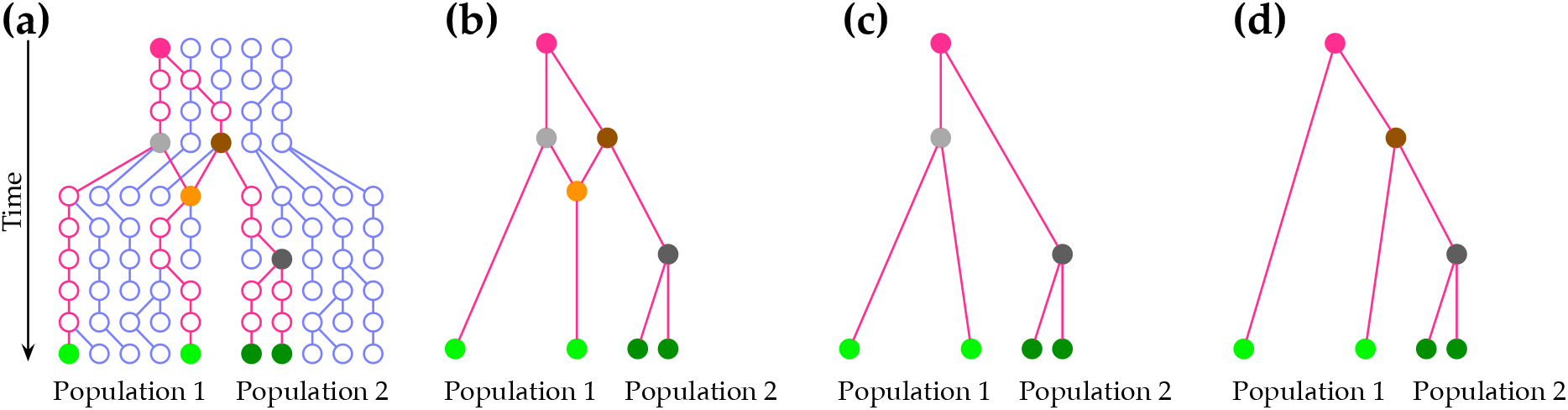
The multispecies coalescent with recombination. (a) The evolutionary history of a sample of four extant individuals in two divergent populations from their MRCA (solid red circle). The recombination node (solid orange circle) results in an ancestral recombination graph (ARG), shown in (b). (c) The genealogy of genomic regions that trace their evolution back from the recombination node to the gray ancestral node. (d) The genealogy of genomic regions that trace their evolution back from the recombination node to the brown ancestral node.

As McVean and Cardin (2005) noted, the coalescent with recombination is very difficult to estimate likelihoods under due to, at least, three important issues: (1) the state-space of recombining genealogies (also known as *ancestral recombination graphs*, or ARGs, illustrated in Fig. 1b) is huge; (2) the data are generally not very informative about the actual ARG; and (3) likelihood estimation is a missing-data problem with highly redundant augmentation.

The consequence of these issues is seen in BACTER (Vaughan et al., 2017), a Bayesian method that uses MCMC to infer the ARG posterior distribution down to the coalescent and recombination times and genomic boundaries of recombinant segments. While this kind of joint inference yields the most detailed posterior distribution, calculating the likelihood of an ARG scales poorly as the number of recombinations increases, and BACTER is limited to analysis of an unstructured population (Vaughan et al., 2017).

Coalescent inference can be scaled up using multilocus methods that assume each locus is spaced far enough apart so that there is effectively no linkage between loci, and each locus is short enough so that no recombination has occurred within it. However, the posterior distribution of local genealogies will be diffuse and incomplete, and the assumption of no recombination within loci has been called into question (Springer and Gatesy, 2016).

To strike a balance between the scalability of multilocus methods and the power of ARG inference, the coalescent with recombination can be approximated as a sequential Markovian process operating across the genome, rather than operating in time along the branches of the phylogeny (Hein, Schierup, and Wiuf, 2005). Using this view, the coalHMM (for “coalescent hidden Markov Model”) was introduced (Hobolth et al., 2007). In this model, an HMM is built such that every coalescent history (gene history) given the species tree is modeled by a state, the transition probabilities are derived based on the recombination rate and the given genealogies, and the emission probabilities are given by the likelihood of the gene trees (Felsenstein, 1981). Fig. 1c,d show two gene histories that are embedded inside the ARG shown in Fig. 1b.

In the work of (Hobolth et al., 2007), the authors determined the transition probabilities by careful inspection of recombination scenarios given the species tree. Later, Dutheil et al. (2009) provided a detailed mathematical derivations under the coalescent with recombination of the model of Hobolth et al. (2007). Such a manual approach to deriving transition probabilities has limited coalHMM-based inference of evolutionary parameters to three genomes. Several later works have attempted to automatically create coalHMMs for various demographic scenarios. For example, Mailund, Halager, and Westergaard (2012) used colored Petri nets to represent genetic models and gave an algorithm for translating such models into coalHMMs. However, there is no software associated with the method. Cheng and Mailund (2020) developed the Jocx tool for ancestral population genomics inference based on pairwise coalescent hidden Markov models, but Jocx currently only supports a limited number of demographic models.

In this work, we present a new method, VICAR (**V**ariational **I**nference under the **C**o**A**lescent with **R**ecombination) for approximate inference under the multispecies coalescent with recombination. VICAR consists of two novel components. First, it implements a simulation-based technique for automatically deriving a coalHMM on which likelihood calculations are done efficiently. More specifically, the likelihood of a candidate model (species tree, divergence times, and population sizes) is computed by simulating data under the coalescent with recombination using the candidate model, using this data to automatically construct an HMM, and then computing the likelihood by means of the Forward algorithm (Chang and Hancock, 1966; Baum et al., 1970; Baum, 1972). This way the method is able to automatically generate coalHMMs and learn their parameters for arbitrarily complex demographic scenarios. By learning coalHMMs from simulation, VICAR also infers local genealogies for arbitrary demographic models. Second, the parameter estimation in VICAR consists of a novel application of variational inference for Bayesian inference of demographic parameters using this approximate likelihood (the species tree topology is assumed to be known and fixed).

We demonstrate the utility and accuracy of VICAR on both simulated and biological data, and compare it to diCal2 (Steinrücken et al., 2019), which is the current state-of-the-art coalescent hidden Markov model algorithm for inferring population histories. Furthermore, we discuss and provide preliminary results for how to scale the method to larger numbers of taxa using a divide-and-conquer approach. The automated nature of our method provides a step towards wider applicability of the coalescent with recombination.

## Results

### Overview of VICAR

Given the topology of a species tree, we seek to estimate its continuous parameters from the genomic data under the (multispecies) coalescent with recombination. As we stated above, maximum likelihood estimation of the topology’s parameters under the exact complex model of the coalescent with recombination is intractable. We introduce a novel variational Bayesian method, VICAR, for accomplishing this estimation by using a simulation-based likelihood kernel. The kernel automatically derives an empirical, simulation-based coalHMM and performs the likelihood computations on the HMM, which can be done in polynomial time in the number of states (Durbin et al., 1998). This automated procedure of generating a coalHMM and computing the likelihood obviates the need for theoretical derivations based on the coalescent theory for every evolutionary scenario as in most previous coalHMM methods, and thus can be applied to infer parameters of any demographic model. We now give a high-level description of how VICAR works, and then give the full details in Methods.

Let Ψ be a species tree on set 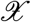 of taxa, **N** be a vector of the effective population sizes associated with Ψ’s internal and root branches, and **T** be a vector of the divergence times of the internal nodes of Ψ. Let Θ = [**N**; **T**]. We fix the hyper-parameters of the prior distributions on the parameters Θ. We also assume a fixed mutation rate *μ* and recombination rate *ρ*. VICAR assumes one sequence from each extant population so the posterior distribution of tip branch population sizes will be identical to the prior and, hence, not estimated by the method. VICAR seeks to estimate the posterior distribution over the model parameters, given by

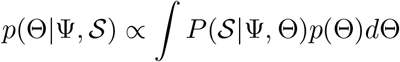

where 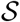 is a genomic sequence alignment that is assumed to have evolved under the coalescent with recombination and a model of sequence evolution. VICAR estimates 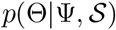 via a novel method of approximating the likelihood 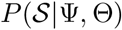 and a novel application of variational inference to estimate the integral.

For approximating the likelihood, VICAR simulates a set of genealogies under the coalescent with recombination and molecular sequences down these genealogies given a model of evolution. Using the simulated data, VICAR constructs an empirical HMM and applies an existing polynomial-time algorithm to compute the likelihood of the HMM. For approximating the posterior distribution, we introduce a novel application of variational inference to this domain. VICAR produces estimates of the parameter values Θ along with measures of confidence.

### Simulation Study on a Human-Chimp-Gorilla Scenario

In this section we demonstrate the performance of VICAR on simulated data and compare it to that of diCal2 (Steinrücken et al., 2019), which is the current state-of-the-art coalescent hidden Markov model algorithm for inferring population histories. diCal2 is based on the sequentially Markov conditional sampling distribution framework (Paul, Steinrücken, and Song, 2011; Steinrücken, Paul, and Song, 2013; Sheehan, Harris, and Song, 2013) with a combination of expectation-maximization (EM) and genetic algorithm to infer a maximum likelihood point estimate, which differs from our Bayesian approach.

We used the program `msprime` (Kelleher, Etheridge, and McVean, 2016) as the simulator for coalescent with recombination process, and `INDELible` (Fletcher and Yang, 2009) as the sequence evolution simulator. We simulated 100 data sets with 500,000 sites each, intended to resemble human-chimp-gorilla (HCG). We refer to the three extant species as human (H), chimp (C) and gorilla (G), and the ancestral species as the human-chimp ancestor (HC) and the human-chimp-gorilla ancestor (HCG). The simulation setup consists of two steps. In the first step, we took the demographic parameters of a species tree and simulated under the coalescent with recombination process. This step gave us a set of segments of the sequence, where each segment had a corresponding coalescent tree. The second step used standard evolutionary simulators to generate sequence alignments at each segment at the given substitution rate under the coalescent tree at that segment. The result of the simulation was a sequence alignment for the set of taxa, where different sites in the alignment had potentially different genealogies. The continuous parameters used in simulation are population sizes *N_HC_* = *N_HCG_* = 40,000, *N_H_* = *N_C_* = *N_G_* = 30,000, speciation time *T_HC_* = 160,000 generations (or 4 Myr assuming a generation time of 25 years), speciation time *T_HCG_* = 220, 000 generations (or 5.5 Myr assuming a generation time of 25 years). The recombination rate is *r* = 1.5 × 10^−8^ per site per generation, corresponding to a genetic recombination frequency of 1.5 cM per Mb. The mutation rate is 2.5 × 10^−8^ per site per generation, corresponding to 0.1% change per million years assuming a generation time of 25 years. The parameters are the same as used in the other two human-chimp-gorilla simulation studies of coalescent HMM (Hobolth et al., 2007; Dutheil et al., 2009).

For each data set, VICAR was used to find the variational posterior of each continuous parameter. The configuration for constructing simulation-based coalHMM as in Algorithm 1 is *nb* = 2 and `-r` = 1000. The simulation length *ℓ* is determined automatically using a scaling formula depending on the `-r` setting, as described below. For stochastic variational inference search, we used 50 samples per iteration with 200 iterations. We used uniform prior on node heights and gamma prior on population sizes. For diCal2, we used 70 samples for each generation of the genetic algorithm, each optimized for 4 EM iterations, and 6 parents for the next generation. We ran the genetic algorithm for 5 generations (the same setting as in (Steinrücken et al., 2019)), and reported the maximum likelihood parameters for each data set.

Fig. 2 shows the maximum a posteriori (MAP) estimates obtained by VICAR and maximum likelihood estimates (MLE) obtained by diCal2 as a violin plot. All parameters are estimated by VICAR with very high accuracy and little variance. Generally, population sizes are estimated with larger variance than node heights, which is true for both methods. VICAR produces more accurate estimates than diCal2 for all four parameters. Furthermore, diCal2 significantly underestimates the ancestral population size of human-chimp ancestor, while VICAR infers the parameter with much higher accuracy and less variability.

**Figure 2:**
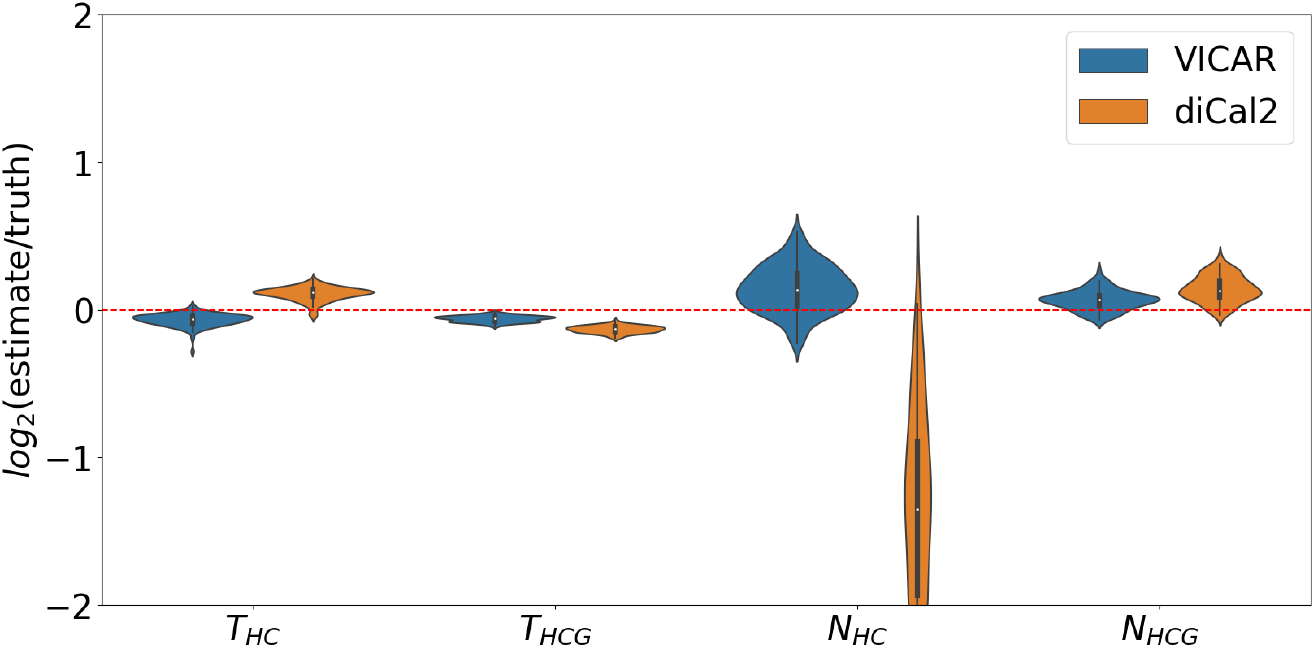
Accuracy results of VICAR and diCal2 on simulated human-chimp-gorilla data sets. The violin plot shows the base-2 logarithm of the relative error (estimate/truth) for the analysis of 100 data sets by VICAR (in blue) and diCal2 (in orange). A value of 0 (the red dashed line) represents an exact estimate.

All the experiments reported above were run on a Macbook Pro with 2.4GHz Intel Core i5 CPU. On average, the run-time of VICAR is about 10.08 hours for a data set. Building the coalHMM by simulation takes 3.93 hours, and computing the likelihood using the Forward algorithm takes 6.15 hours. diCal2 takes less time, with only 0.8 hours per data set. This can be partially explained by the fact that diCal2 makes further approximations to make the HMM more efficient, and incorporates optimizations for speeding up the Forward algorithm, while our current implementation of VICAR does not optimize the HMM and applies a vanilla implementation of the Forward algorithm. To study how the two methods perform given comparable computational cost, we assessed the performance of VICAR based on the partial results obtained after the first 20 iterations of variational inference search, which takes about one hour to run (comparable to the running time of diCal2). As the results in Fig. 3 show, VICAR still achieves similar or better estimates for three of the four parameters, and much more accurate estimates for *N_HC_*.

**Figure 3:**
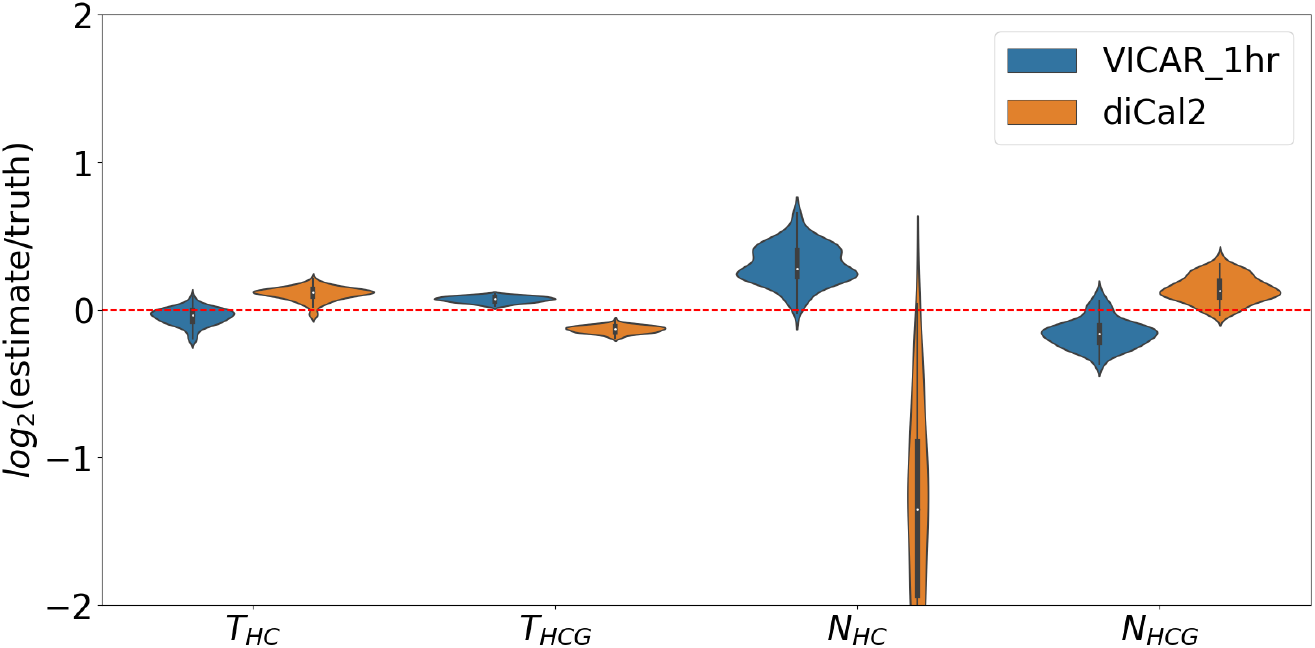
Accuracy results of VICAR (blue) and diCal2 (orange) on simulated human-chimp-gorilla data sets given comparable computational cost. Here, VICAR was run for only 20 iterations in the variational inference step.

For a more general view of VICAR’s performance as a function of the number of VI iterations, Fig. 4 shows the convergence plot of VICAR on one of the data sets. As the figure shows, the likelihood increases rapidly in the first few iterations, and converges after only about 50 iterations.

**Figure 4:**
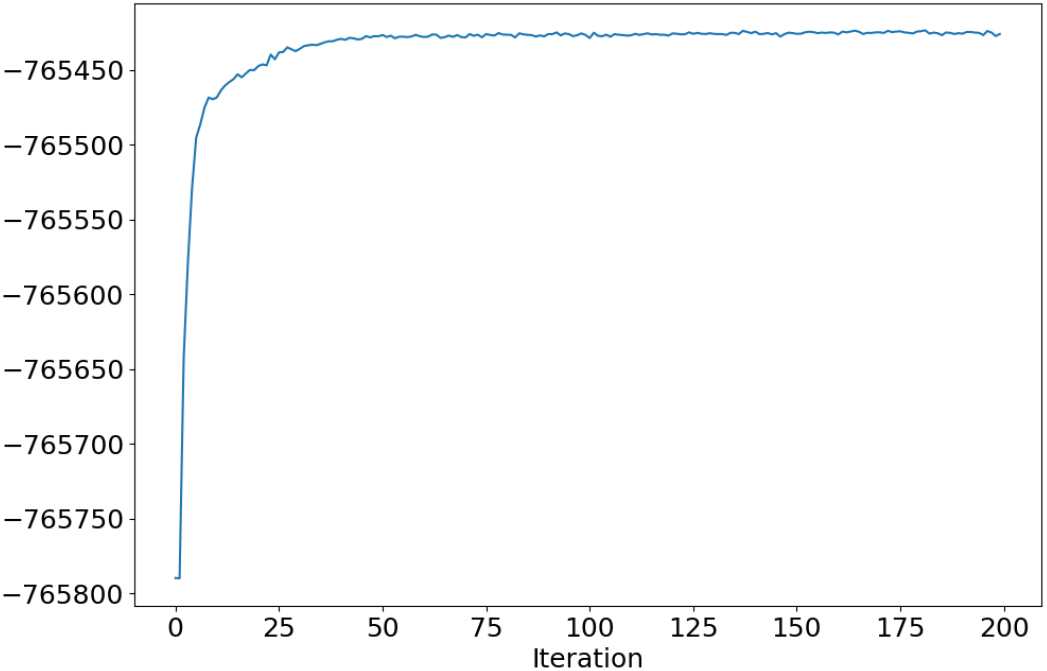
Convergence of VICAR. The x axis is the number of iterations and the y axis is the log likelihood of the current estimation.

Another advantage of our method is that it is a Bayesian approach, with the ability to specify priors and provide posterior support for parameters. Once VICAR converges, the uncertainty associated with the inference is quantified by the Bayesian posterior, while for maximum likelihood approaches such as diCal2, running the algorithm multiple times for bootstrapping is required to infer confidence intervals, which increases the actual running time.

Fig. 5 shows, for each parameter, the number of inferences where the true value is within the 95% credible interval of the estimation. As the figure shows, the true values of all parameters fall within the 95% credible interval most of the time.

**Figure 5:**
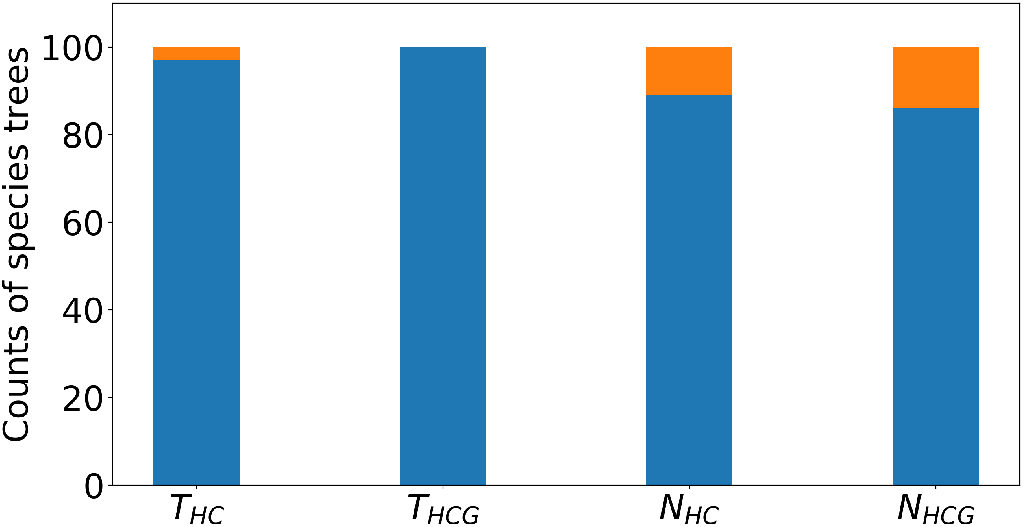
Accuracy of variational posteriors. For each parameter, the number of inferences with the true value within 95% credible interval of the estimation (blue) and outside 95% credible interval of the estimation (orange) are shown.

### Analysis of a Biological Data Set

We reanalyzed the empirical human-chimp-gorilla sequences from Hobolth et al. (2007) for comparison with previous models. We reanalyzed target 106 (Chromosome 20) of Hobolth et al. (2007) using VICAR and diCal2 and compared the results with those reported in Hobolth et al. (2007) and Dutheil et al. (2009) on the same data. The VICAR settings were the same as in the simulation study above. We used a recombination rate *r* = 2 × 10^−9^ per site per generation, and a mutation rate *μ* = 2.35 × 10^−8^ per site per generation, as they were estimated from pedigree data and reported in Dutheil et al. (2009) for this target. The result is shown in Fig. 6. Generally, VICAR infers comparable results to those reported in Hobolth et al. (2007) and Dutheil et al. (2009), whereas diCal2 yields farther estimates for the two population sizes. For *T_HC_*, all four methods infer about the same value. For *T_HCG_*, VICAR’s estimate is closer to that of Dutheil et al. (2009) than the other two.

**Figure 6:**
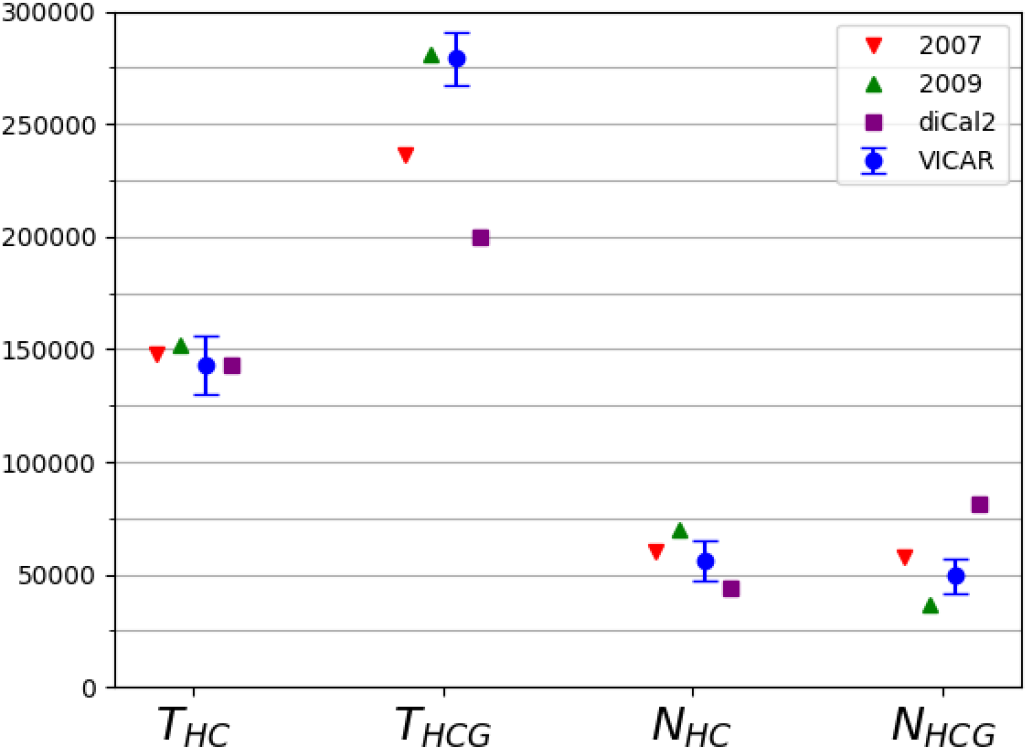
Inference results on target 106. The factorized normal variational posterior distribution of each parameter inferred by VICAR is shown in blue. The red triangle shows the maximum likelihood solution obtained by Hobolth et al. (2007). The green triangle shows the maximum likelihood solution obtained by Dutheil et al. (2009). For the 2009 model, we took the result after bias correction. The purple square shows the solution obtained by diCal2, and the blue circle is the VICAR solution. Given that VICAR employs Bayesian inference, it provides confidence measures; shown are standard deviations of the Gaussian posteriors.

### Local Genealogy Inference

Other than inferring continuous parameters, an important capability of VICAR is the inference of the local genealogy of each site along the genome. Since the hidden states of a coalescent hidden Markov model are coalescent histories (genealogies), local genealogy inference can be done by posterior decoding of the HMM along the sequence data, which gives us the posterior probability of each genealogy at each site. In this section, we study the performance of our simulation-based HMM in terms of local genealogy inference.

We used the same 100 simulated data sets as in the simulation study above. Since we used `msprime` to simulate under the coalescent with recombination process when generating data, we have the true coalescent tree of each site. We used *nb* = 2 and `-r` = 1000 to build our HMM by simulation.

For the human-chimp-gorilla species tree, there are four types of genealogies: HC1, HC2, HG, and CG, as shown in Fig. 14a. Since we fine-grained each branch into two sub-branches, our HMM has a higher granularity than four genealogies. The total number of states in our HMM is actually 13. However, for the purpose of local genealogy inference, we only consider the four basic types as they are the most meaningful categorization for determining the shared ancestry of molecular characters or traits. Therefore, after posterior decoding on the 13-state HMM, we merged hidden states of the same type together and took the type with the highest posterior probability as the inferred genealogy at each site. We also discretized the true coalescent tree at each site into one of the four genealogies. We then compared the inferred genealogy with the true one. Table 1 shows the confusion matrix of the classification task, as well as the precision and recall measures for each type of genealogy. Fig. 7 shows a graphical comparison of the posterior probabilities of each genealogy at each site with the true genealogy along the sequence from a segment of 100,000 sites from one of the data sets.

**Figure 7:**
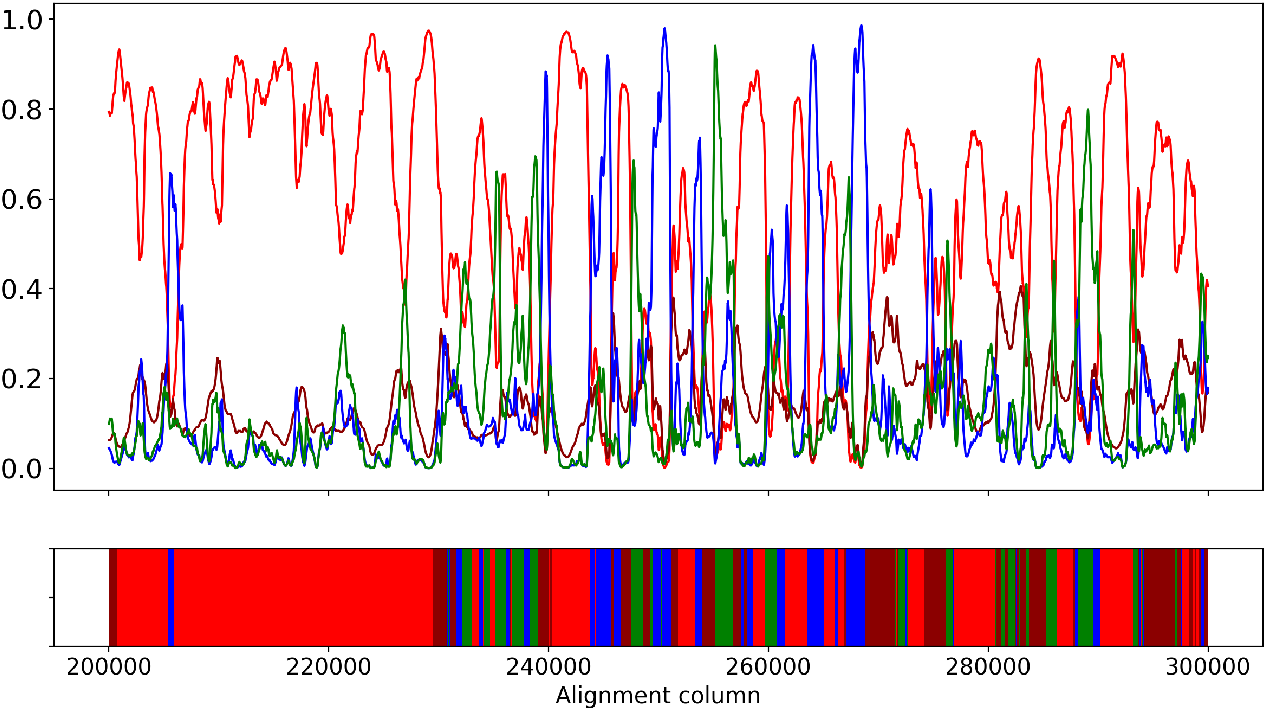
True genealogy and posterior distribution along the sequence. The upper panel shows posterior probability of each genealogy at each site. The lower panel shows the true genealogy of each site. Coloring corresponds to different genealogies: genealogy HC1 is in red, HC2 is in dark red, HG is in blue, and CG is in green.

**Table 1:** Classification accuracy of local genealogies on 100 simulated data sets. (a) A confusion matrix of the genealogy classification task based on posterior decoding. Sums over rows give the frequencies of true genealogies, and sums over columns give the frequencies of inferred genealogies. The diagonal corresponds to correctly inferred cases. (b) The precision measure for each genealogy, as defined by the number of true positives over the number of positives. (c) The recall measure for each genealogy, as defined by the number of true positives over the true number of sites with that genealogy.

| Inferred genealogy<br>True genealogy | (a) Posterior Decoding |  |  |  | (b) Precision | (c) Recall |
| --- | --- | --- | --- | --- | --- | --- |
|  | HC1 | HC2 | HG | CG |  |  |
| HC1 | 47.56% | 0.52% | 2.23% | 2.32% | 66.73% | 90.38% |
| HC2 | 11.32% | 0.70% | 1.89% | 1.93% | 38.44% | 4.44% |
| HG | 6.17% | 0.30% | 7.12% | 2.02% | 53.27% | 45.60% |
| CG | 6.21% | 0.31% | 2.12% | 7.26% | 53.67% | 45.64% |

The results show that the accuracy of local genealogy inference is very good: 90% of true HC1 sites are inferred to have HC1 genealogy (Table 1). This number is 46% for HG and CG, with an overall classification accuracy of 63%. This number is significantly higher than random expectation (Dutheil et al., 2009). We observe the same good performance in Fig. 7, where there is a good correspondence between true genealogy and posterior distribution. However, note that the recall measure of HC2 is only 4.44%, meaning only 4.44% of all true HC2 genealogies are actually estimated to be HC2. Dutheil et al. (2009) reported the same poor performance on HC2. Many sites with HC2 as true genealogy are assigned to another type, mostly HC1 (Dutheil et al., 2009). This is likely a model artifact of the HMM approximation to the coalescent with recombination process. We already know that the HMM approximation would underestimate the recombination rate (Dutheil et al., 2009; Mailund et al., 2011), which means it would underestimate state transitions, leading to a global underestimation of incomplete lineage sorting. For most of the true HC2 sites, it is unsurprising that these sites are misclassified as HC1, since the stationary frequency of HC1 is so much higher and given that the site patterns of true HC1 sites and true HC2 sites should be similar.

### Relationship Between Inference Accuracy, Number of Sub-branches, Simulation Length, and Branch Lengths

The accuracy of inferences depends on the quality of the approximate likelihood, which in turn depends on two aspects: the accuracy of the simulation-based coalHMM approximation of the coalescent with recombination process, and the quality of the trained coalHMM itself. The accuracy of the coalHMM approximation of the coalescent with recombination process is determined by the refinement of coalHMM state space, *i.e*., the number of sub-branches on each branch of the species tree when building the HMM. If the number of sub-branches is small, the resulting coalHMM has a state space of coarse coalescent histories not enough to capture the detailed coalescent distribution, leading to biased likelihood (Mailund et al., 2011; Dutheil et al., 2009). The quality of the coalHMM itself (*i.e*., the quality of the transition matrix) is determined by the length of simulation used to derive the HMM since our coalHMM is trained directly from labeled sequence data. The more sub-branches we use to refine a branch, the more accurate approximation we obtain, but more subbranches incur a larger state space, necessitating a longer simulation length in order to train a high-quality transition matrix. If we use a large number of sub-branches but a small simulation length, the resultant HMM will be unreliable because of limited training data. Moreover, depending on the branch length, we may not need a large number of sub-branches to approximate the coalescent process on that branch, but a number of sub-branches too small may introduce bias. In this section, we study the relationship between accuracy of inference result, number of sub-branches used to refine HMM state space, length of simulation used to build HMM, and branch length of the species tree we do inference on. We derive some empirical suggestions on what number of sub-branches and simulation length to use for any specific inference problem.

There are two hyper-parameters controlling the HMM building process in Algorithm 1: *nb*, the number of sub-branches on each branch of the species tree, and *l*, the length of simulated HMM training data. In our implementation, they are user-defined inputs `NUM_BIN` and `CROSS_OVER_RATE. NUM_BIN` is the number of sub-branches used to approximate each branch of the species tree. `CROSS_OVER_RATE` is the `-r` switch of `ms` (Hudson, 2002), which is defined as 4*N*_0_*r*, where *N*_0_ is a customized population size and *r* is the probability of recombination per generation between the ends of the locus being simulated, which is the probability of recombination per site per generation times the number of sites to simulate. Our implementation assumes a value of 10,000 for *N*_0_, and it can be changed according to the scale of population sizes. For example, if the recombination rate is 1.5 × 10^−7^/site/generation and we seek to simulate 500000 sites, we use 4 × 10000 × 1.5 × 10^−7^ × 500000 = 3000 as the value for the `-r` option.

We simulated sequence alignments under three different evolutionary scenarios, the difference between which is the internal branch length. All three scenarios have the same three-taxon tree topology ((*A, B*),*C*);. Internal node height *T_AB_* is fixed at 100,000 generations. The root node height varies across scenarios. Scenario 1 has root node height 150,000 generations. Scenario 2 has root node height 200,000 generations. Scenario 3 has root node height 400,000 generations. Population sizes of all branches are fixed at 40,000. Hence, the branch length of the internal branch for scenarios 1, 2, and 3 in coalescent units are 0.625, 1.25, and 3.75, respectively. The recombination rate is *r* = 1.5 × 10^−7^/site/generation. The mutation rate is 1.25 × 10^−6^/site/generation. The length of the sequence is 100,000 bp.

We inferred the continuous parameters of each scenario under various configurations of the hyperparameters. The number of sub-branches explored were 1, 2, 3, 4, and 5. The values for the `-r` parameter explored were 500, 1000, 3000, and 5000, which correspond to simulation lengths of 83333, 166666, 500000, and 833333, respectively. For each combination of number of sub-branches and `-r` parameter value (simulation length), we conducted inference on each scenario using the combination for coalHMM construction to find the MAP solution, and inspected the accuracy. In total, we conducted 3 × 4 × 5 = 60 inferences.

Figs. 8, 9, and 10 show inference results for scenario 1, 2, and 3, respectively. The message is clearest in Fig. 10. Looking at each individual plot, for a fixed simulation length, increasing the number of sub-branches increases the accuracy of inference, until it flattens out. But Fig. 8 suggests that the accuracy does not stay on a plateau after reaching a certain number of sub-branches. Rather, if the simulation length is not long enough, increasing the number of sub-branches might decrease inference accuracy. The reason is that the transition rate matrix of a large state space cannot be sufficiently trained from a short length of simulation. Each column of the plots show that generally, for a fixed number of sub-branches, increasing the simulation length increases the accuracy, but the gain is smaller than increasing the number of sub-branches. Simulation length only determines how close is the transition matrix of a coalHMM trained from simulated data to the true transition rate matrix calculated from a strict mathematical model. However, the bias in approximate likelihood comes from a restricted state space, not a poor transition matrix. If the state space is restricted, increasing simulation length does not solve the bias problem because the bias still exists when the transition rate matrix is analytically derived (Mailund et al., 2011; Dutheil et al., 2009). Put simply, increasing the simulation length is not going to address any biases resulting from discretization the state space of coalescent histories, or from using a Markov chain to approximate the ARG.

**Figure 8:**
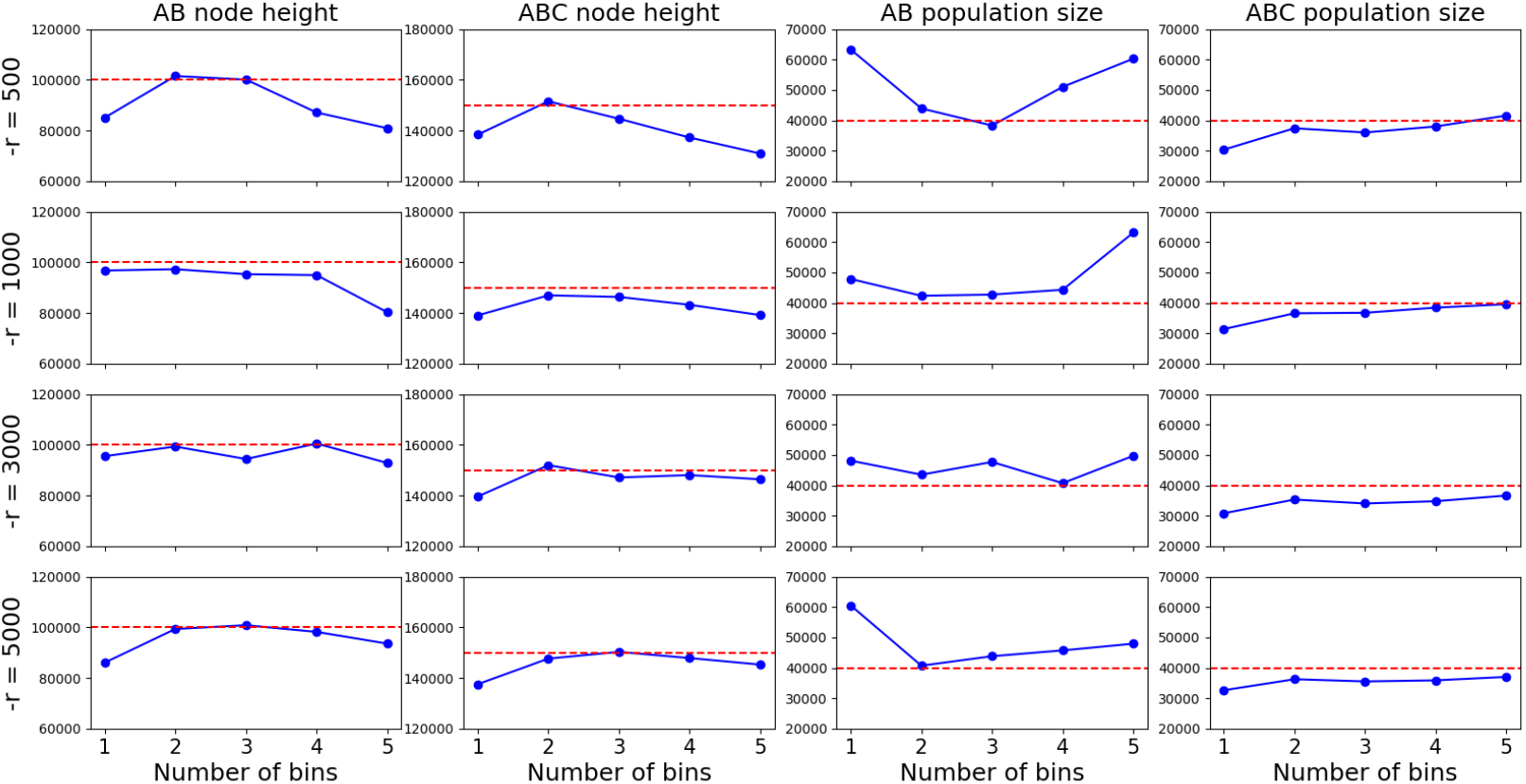
Results on scenario 1 (internal branch length 0.625). Dashed red lines are true values. Blue lines are inferred MAP values. Rows correspond to different simulation lengths. Columns corresponds to different continuous parameters. x-axes are number of sub-branches ranging from 1 to 5.

**Figure 9:**
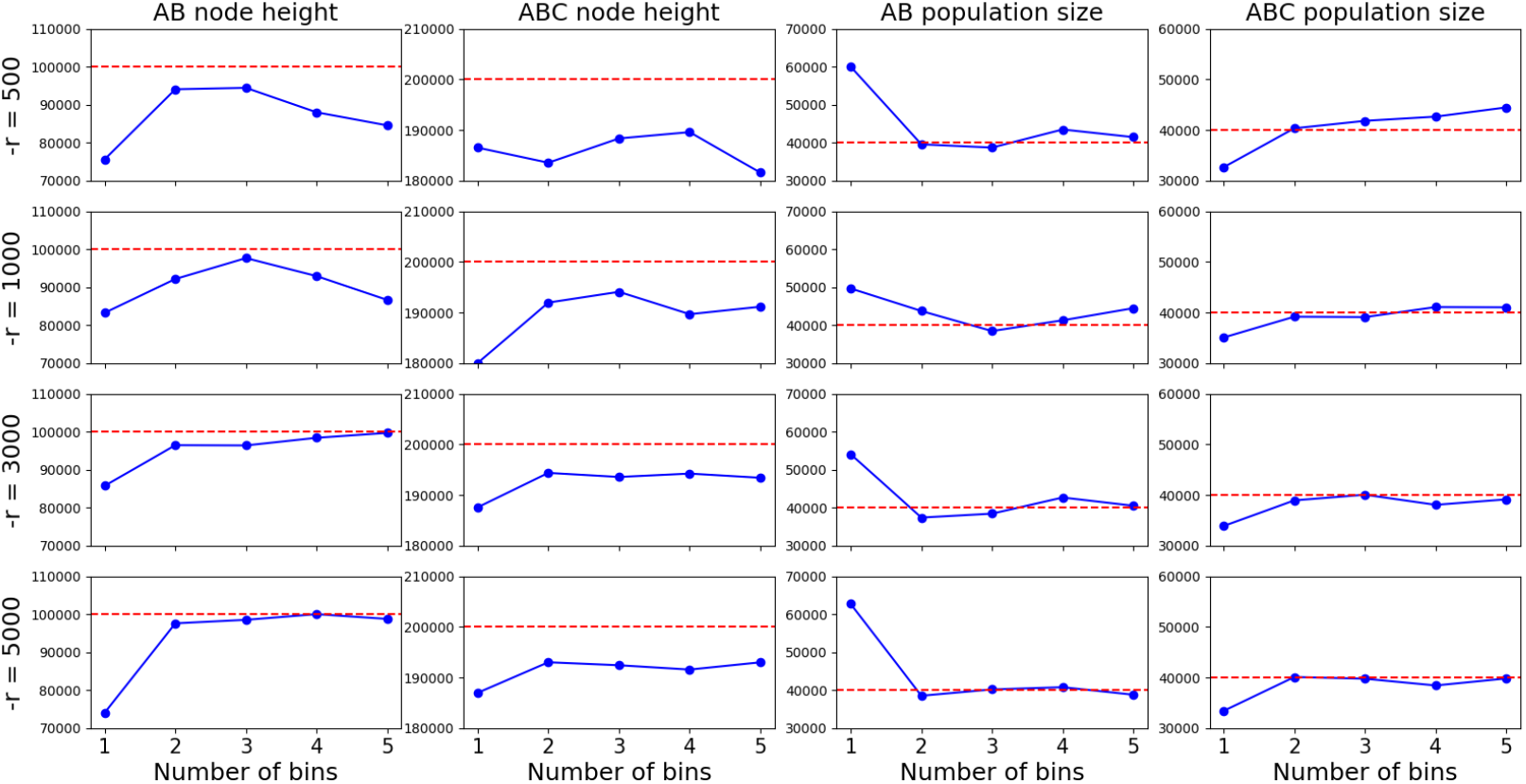
Results on scenario 2 (internal branch length 1.25). Dashed red lines are true values. Blue lines are inferred MAP values. Rows correspond to different simulation lengths. Columns corresponds to different continuous parameters. x-axes are number of sub-branches ranging from 1 to 5.

**Figure 10:**
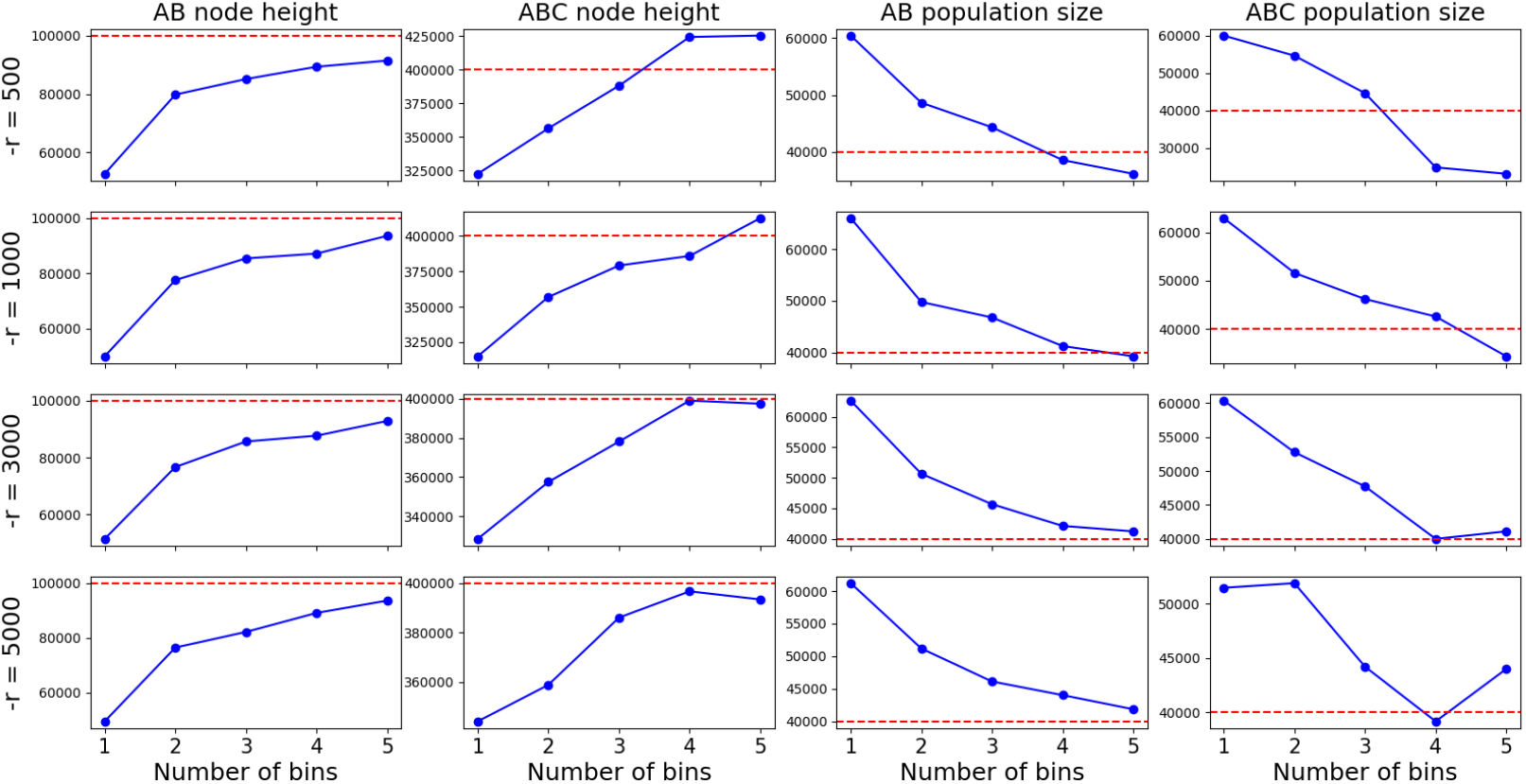
Results on scenario 3 (internal branch length 3.75). Dashed red lines are true values. Blue lines are inferred MAP values. Rows correspond to different simulation lengths. Columns corresponds to different continuous parameters. x-axes are number of sub-branches ranging from 1 to 5.

The appropriate number of sub-branches and simulation length to use depends on the internal branch length of the species tree. For example, using two sub-branches and `-r` = 1000 infers a very good result on scenario 1, but does not work well on scenario 3. Based on the plot, for a short internal branch (about one coalescent unit), two or three sub-branches with a `-r` of around 1000 are sufficient. For a larger branch length (one to three coalescent units), three or four sub-branches with a `-r` of about 3000 would suffice. For branches longer than three coalescent units, more than four sub-branches and a `-r` value higher than 3000 would be needed.

Running time is also a consideration when choosing hyper-parameters. We now study the impact of these hyper-parameters on running time. Almost all the time taken by the inference attributes to Algorithm 1. The algorithm has two time-consuming sub-procedures: building the coalHMM by simulation, and calculating the likelihood by the Forward algorithm. The running times of the two sub-procedures depend on the number of sub-branches, the length of the simulation, and the scale of the species tree. Fig. 11 shows the relationship between time taken by the Forward algorithm and the number of sub-branches for refining the species tree, while keeping all other variables fixed, when evaluating the approximate likelihood of one model. Clearly, the number of sub-branches has a huge impact on the running time, as the Forward algorithm time significantly increases with the number of sub-branches. The reason is that the number of hidden states of the resulting coalHMM is a high-degree polynomial in the number of sub-branches, which results in large increase in the Forward algorithm running time since it is quadratic in the HMM size. Fig. 12 shows the relationship between the HMM building time, the Forward algorithm running time, and the length of simulation used to build coalHMM, while keeping all other variables fixed. We observe that the time taken for building the HMM grows with simulation length, which has to do with the complexity of the underlying simulator, in our case `msprime`. The Forward algorithm running time remains roughly unchanged since the number of hidden states of the coalHMM does not depend on simulation length.

**Figure 11:**
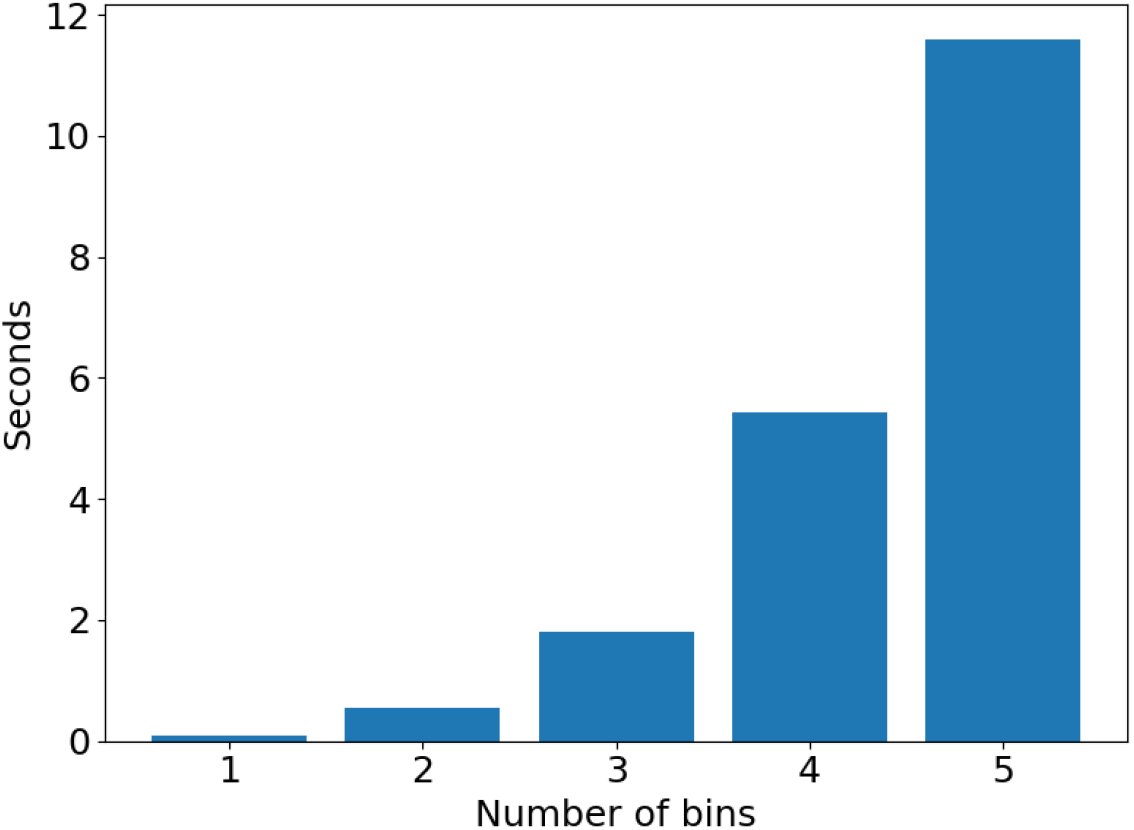
Running time of the Forward algorithm. The actual running time taken by the Forward algorithm when calculating the approximate likelihood of Scenario 3, with different *nb* parameter values in Algorithm 1.

**Figure 12:**
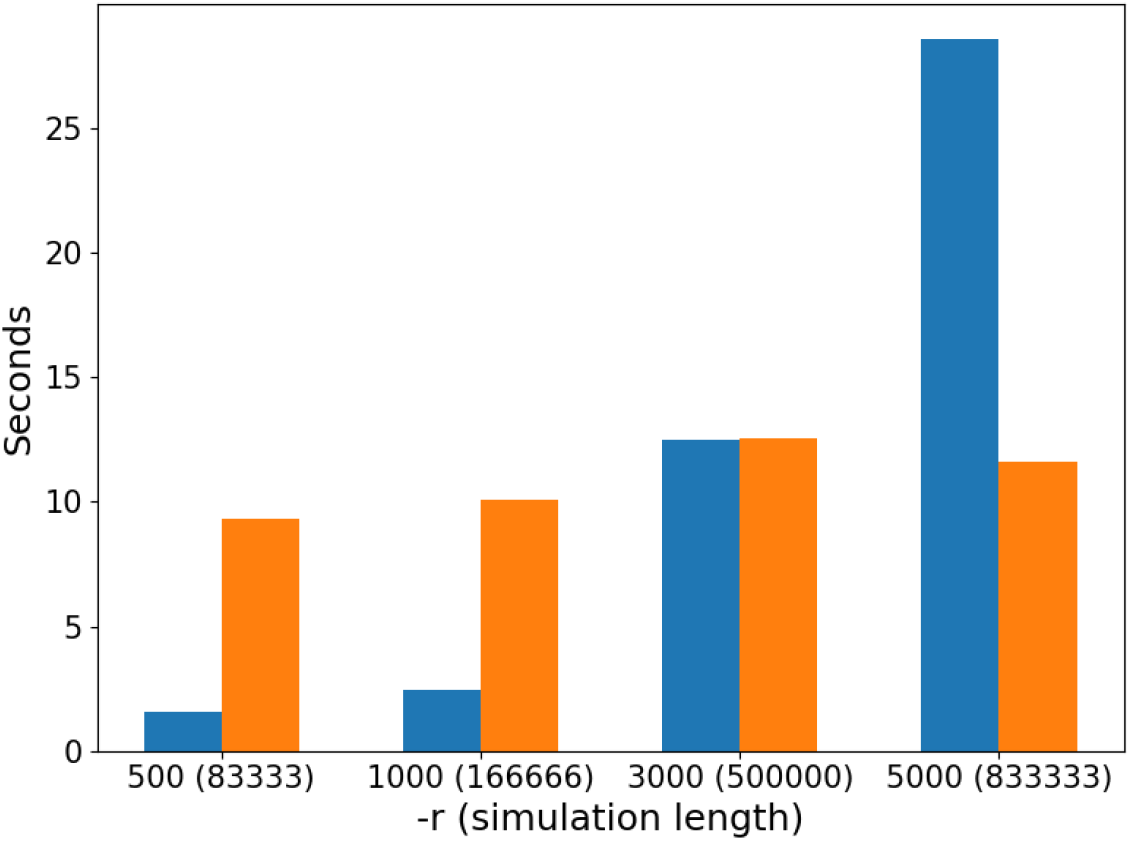
Running times of building the HMM and the Forward algorithm. The actual running times taken by building the HMM by simulation (blue bars) and running the Forward algorithm (orange bars) when calculating the approximate likelihood of Scenario 3, with different `-r` parameter values (the *l* parameter in Algorithm 1).

Fig. 13 shows the relationship between the HMM building running time, the Forward algorithm running time, and the branch length of the species tree on which the approximate likelihood is being evaluated. As before, the time taken for building the HMM grows, though not exponentially, with the simulation length, while the Forward algorithm running time remains unchanged. These results highlight the need for developing scalable simulators under the coalescent with recombination, which we identify as a future research direction.

**Figure 13:**
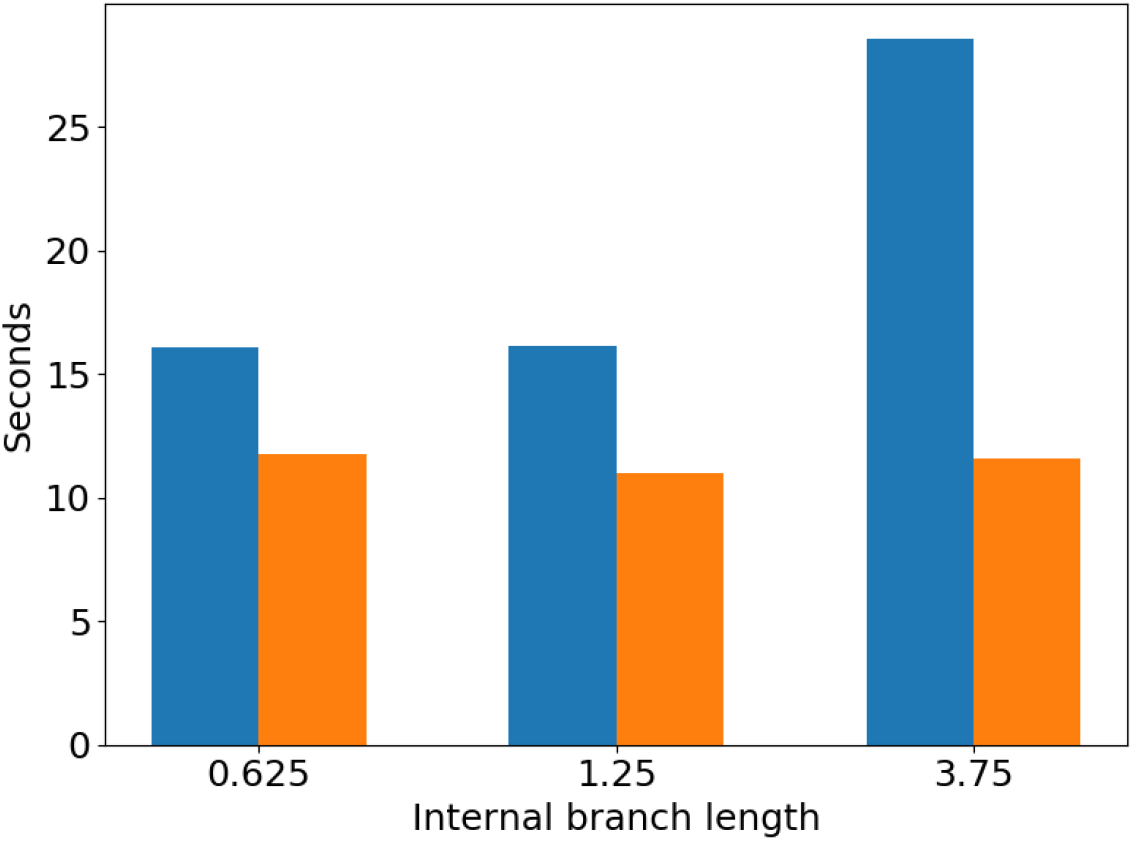
Running times of building the HMM and the Forward algorithm. The actual running times taken by building the HMM by simulation (blue bars) and running the Forward algorithm (orange bars) when calculating the approximate likelihood of Scenarios 1, 2, and 3.

### Simulation Study on a Four-taxon Data Set

To demonstrate the scalability of VICAR, and the efficiency of the divide-and-conquer approach, we simulated a four-taxon sequence data set on the species tree whose topology is (((A,B),C),D). The demographic parameters are *T_AB_* = 100,000, *T_ABC_* = 160,000, *T_ABCD_* = 450,000. All branches have population size 40,000. The recombination rate is r = 1.5 × 10^−7^/site/generation. The mutation rate is 1.25 × 10^−6^/site/generation. The length of the sequence is 200,000 bp. We first ran a full inference on all taxa, and then ran a divide-and-conquer inference to compare the results.

#### Full Inference on Four Taxa

The configuration of the coalHMM likelihood kernel for the full inference is *nb* = 2 and `-r` = 1000. Black box variational inference is set to run for 200 iterations with 50 samples per iteration. We used uniform prior on node heights and gamma prior on population sizes. The inference took 83.09 hours. 7.62 hours were used to build the coalHMM, and 75.46 hours were used to compute likelihood by the Forward algorithm. It is worth pointing out that, due to the increased number of taxa, the number of hidden states of the coalHMM is significantly increased, resulting in a very long running time for the Forward algorithm, which is quadratic in the number of hidden states. This is a major reason why coalHMM methods are limited to a few taxa. The inference results are shown in Table 2. The true values of most of the parameters are within the 95% credible intervals of the estimates. We used two sub-branches per branch, which, based on our results for three taxa, is suboptimal for accuracy, but inference with only two sub-branches took over 80 hours on a Macbook Pro with 2.4 GHz Intel Core i5 CPU. Results using more sub-branches may be more accurate but we stuck with two sub-branches to limit running time.

**Table 2:** Result of full inference on the four-taxon data set. The means and standard deviations of the parameter estimates as well as the true parameter values are shown.

| Parameter | Mean | Standard Deviation | True Value |
| --- | --- | --- | --- |
| $T_{AB}$ | 81770 | 12730 | 100000 |
| $T_{ABC}$ | 145883 | 11568 | 160000 |
| $T_{ABCD}$ | 422377 | 16254 | 450000 |
| $N_{AB}$ | 51343 | 6135 | 40000 |
| $N_{ABC}$ | 37788 | 3430 | 40000 |
| $N_{ABCD}$ | 55340 | 6623 | 40000 |

#### Divide-and-conquer Inference

To reduce the running time and improve the accuracy, we used a divide-and-conquer approach on this data set. As shown in Fig. 15 above, two three-taxon inferences were run to cover all the parameters of the four-taxon tree. The ((A,B),C) subtree covers *T_AB_, T_ABC_*, and *N_AB_*, while the ((A,C),D) subtree covers *T_ABC_, T_ABCD_, N_ABC_*, and *N_ABCD_*. The parameter *T_ABC_* is covered by both data sets.

To infer parameters of the ((A,B),C) tree, we used *nb* = 3 and `-r` = 1000 for building the coalHMM. Black box variational inference settings and prior settings are the same as the full inference. The inference took 5.20 hours. 3.01 hours were used to build the coalHMM, and 2.19 hours were used to compute the likelihood by the Forward algorithm. The parameter estimate results are shown in Table 3.

**Table 3:**
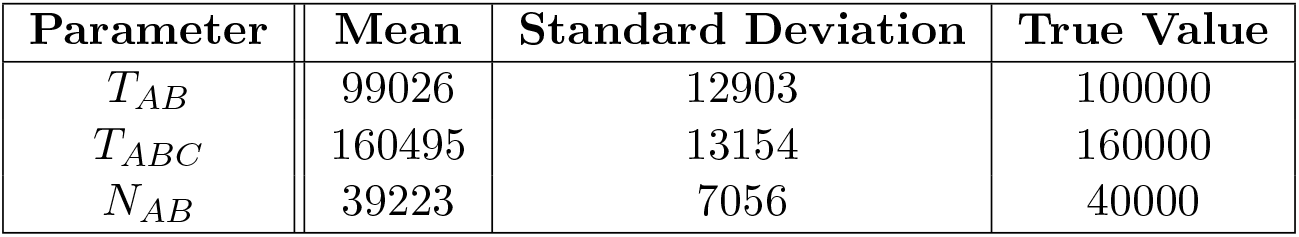
Inference results on the ((A,B),C) tree.

To infer parameters of the ((A,C),D) tree, we used *nb* = 4 and `-r` = 1000 for building the coalHMM. We used one more sub-branch for each branch since the ((A,C),D) tree has a longer internal branch resulting from not sampling taxon B. This further illustrates the flexibility of the divide-and-conquer approach where coalHMM settings can be adjusted according to the specifics of different sub-instances of the problem, saving computational resources overall. The inference took 26.92 hours. 5.41 hours were used to build the coalHMM, and 21.50 hours were used to compute the likelihood by the Forward algorithm. The results are shown in Table 4.

**Table 4:** Inference results on the ((A,C),D) tree.

| Parameter | Mean | Standard Deviation | True Value |
| --- | --- | --- | --- |
| $T_{ABC}$ | 149776 | 11254 | 160000 |
| $T_{ABCD}$ | 445402 | 14242 | 450000 |
| $N_{ABC}$ | 43288 | 4515 | 40000 |
| $N_{ABCD}$ | 42950 | 4531 | 40000 |

As the results in both tables demonstrate, all the parameters are recovered with good accuracy in both subtrees, a significant improvement on the direct full inference in Table 2. An another significant improvement is the running time. While the full inference took 83.09 hours, the two subtree inferences only took 5.20 and 26.92 hours, respectively. Since the two subtree inferences are independent, they can be run in parallel. Hence the divide-and-conquer method reduces the running time of the inference from over 80 hours to a little bit over 25 hours, while achieving a higher accuracy on all parameters. The divide-and-conquer technique is a promising approach towards large-scale population history inference.

## Discussion

Coalescent methods are a fundamental tool of population genetics and increasingly standard in phylogenetics. In particular, the multispecies coalescent (MSC) has emerged as a central model underlying a wide array of methods for inferring species trees that account for the phenomenon of incomplete lineage sorting. However, inference under this model assumes that the data comes from multiple loci such that there is free recombination between loci and no recombination within any locus. This assumption necessitates preprocessing the data carefully before using it as input to the methods. While genomic regions are sampled far enough from each other so as to increase the likelihood of independence among loci, the assumption of no recombination within individual loci is much harder to satisfy when each locus is given by a sequence alignment, as those sequence alignments need to be long enough for phylogenetic signal.

As whole genomes become more affordable and widely available, an alternative approach to inference of evolutionary parameters is to use methods that account for recombination. The (multispecies) coalescent with recombination extends the MSC and allows for modeling the evolution of genomic regions in the presence of coalescent effects as well as recombination. However, inference under this model has thus far proven much more challenging computationally than inference under the MSC. The coalescent hidden Markov model, or coalHMM, framework was introduced for inferences under the MSC with recombination and has offered a promising approach for the analysis of large genomic alignments. In this framework, recombination is viewed as a spatial process operating along the genomic sequence, and an HMM whose states correspond to local genealogies captures the evolutionary.

However, coalHMM methods have been difficult to to generalize beyond a simple three-taxon ultrametric tree. In the work of (Dutheil et al., 2009), the authors conducted detailed mathematical analysis in order to parameterize the transition probabilities of a 4-state coalHMM. Such a manual approach of parameterizing coalHMMs for different species trees is not tenable, and more general inference methods were needed. Most recently, diCal2 (Steinrücken et al., 2019) was introduced for obtaining maximum likelihood estimates of evolutionary parameters under the coalescent (and MSC) with recombination given an arbitrary tree structure of the species or sub-populations. In this work, we presented the method VICAR, which employs a different approach from that of diCal2, for general inference under the MSC with recombination. VICAR uses variational inference for sampling the posterior distribution of the evolutionary parameters, and employs simulations to derive an empirical coalHMM that is amenable to efficient likelihood computations. As VICAR samples the posterior distribution, it naturally provides measures of confidence for the parameter estimates. Furthermore, as VICAR explicitly builds a coalHMM, it can be used in a straightforward manner for obtaining local genealogies for the individual sites (or blocks of sites) in a genomic data set. We demonstrated on simulated data sets that VICAR obtains either comparable or more accurate inferences than diCal2. Furthermore, we discussed how the method can be used in a divide-and-conquer fashion to scale up to larger data sets (in terms of the number of genomes). However, it is important to note that in their current implementations, diCal2 is much more optimized computationally than VICAR (e.g., implementing an algorithm for grouping loci into single large blocks).

Both diCal2 and VICAR assume that the demographic, or evolutionary, structure is known (the tree topology) and focus on estimating the (continuous) evolutionary parameters. Both methods can be coupled with a tree search procedure for a straightforward implementation of evolutionary history inference, including the topology, under the coalescent and MSC with recombination. However, such an implementation could face many of the challenges associated with phylogenetic inference in general due to the discrete nature of the search space as well as the complexity of the likelihood surface and posterior distribution. Furthermore, while in this work we focused on tree-structured models, the approach underlying VICAR is extendible to network structures, thus allowing for modeling gene flow as well, which we identify as a direction for future research. In principle, all kinds of generalizations are possible as long as they can be simulated. Examples would include nonconstant demographic functions such as linear, stepwise or exponential changes in population sizes, and ancient hybridization. Our new approach will be immediately useful to researchers working at the intersection of population genetics and phylogenetics, but also represents an additional step forward in terms of applying coalHMMs to biological systems beyond the relatively simple humanchimp-gorilla tree.

## Methods

### Simulation-based Likelihood Approximation

For a fixed species tree topology Ψ, given a specific Θ and a sequence alignment 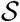, we seek to compute the likelihood of Θ given by 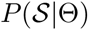. Note that this likelihood marginalizes over the local genealogy at each site. Algorithm 1 gives the procedure for approximate likelihood computation.

**Algorithm 1:**
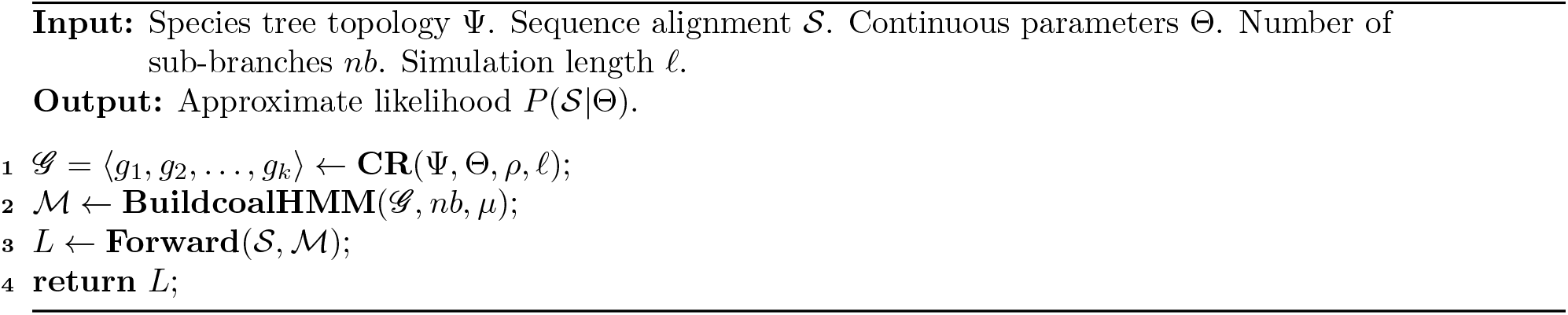
ApproximateLikelihood.

**CR** (for **C**oalescent with **R**ecombination) runs a coalescent-with-recombination simulator to generate a sequence 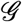 of *k* local genealogies corresponding to *ℓ* sites under the model specified by Ψ and Θ. Here, each of the *k* genealogies correspond to a contiguous genomic region of one or more sites, the genomic regions of the genealogies are pairwise disjoint, and the concatenation of the *k* genomic regions yields a region of *ℓ* sites. Each of the *k* genomic regions is recombination-free, and every two consecutive regions are separated by at least one recombination event. In the implementation, we use `msprime` (Kelleher, Etheridge, and McVean, 2016), a reimplementation of Hudson’s classical `ms` simulator (Hudson, 2002) for efficient coalescent simulations. There is clearly a trade-off between computational requirements and accuracy when setting the value of *ℓ*, which we discuss in the Results section below.

After the sequence 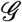 of gene trees is produced, **BuildcoalHMM** empirically builds a coalHMM as follows. In its basic version, **BuildcoalHMM** builds a coalHMM with one state per coalescent history (Degnan and Salter, 2005) given the species tree. For example, for the species tree Ψ in Fig. 14a, the basic coalHMM would have four states corresponding to the four coalescent histories HC1, HC2, HG, and CG. This is precisely the model used in (Hobolth et al., 2007). However, as discussed elsewhere (Dutheil et al., 2009), having one state per coalescent history could result in unidentifiability of some of the parameters. To ameliorate this problem, **BuildcoalHMM** in our method can refine the states further by segmenting branches in the species tree into contiguous non-overlapping sub-branches, and refining individual coalescent histories based on this segmentation. This concept is illustrated in Fig. 14b, where the internal branch separating (H,C) from the root of the tree is segmented into three sub-branches. Now, coalescent history HC1 of Fig. 14a is refined into three coalescent histories, HC1.1, HC1.2 and HC1.3, each corresponding to a unique mapping of the coalescent history of h and c to a sub-branch. The number of sub-branches is controlled by the parameter *nb* in Algorithm 1. We explore the impact of *nb* on the accuracy and computational requirements in the Results section below. Finally, the transition probabilities are derived empirically from the simulated coalescent histories, estimating the rate of transition from one history to another by simple counting of the number of transitions in the simulation. By empirically constructing the coalHMM from simulated sequence, we also potentially reduce the state space explosion problem of dealing with many populations (Cheng and Mailund, 2020; Mailund, Halager, and Westergaard, 2012). Only coalescent histories that appear in the simulation are taken into account, and states that are not simulated are omitted without affecting the accuracy of the likelihood, as we demonstrate below. The emission probabilities for each state at each alignment column of 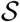 are computed by Felsenstein’s pruning algorithm (Felsenstein, 1981). In the implementation, we use the BEAGLE library (Ayres et al., 2012) for efficient implementation of Felsenstein’s algorithm.

**Figure 14:**
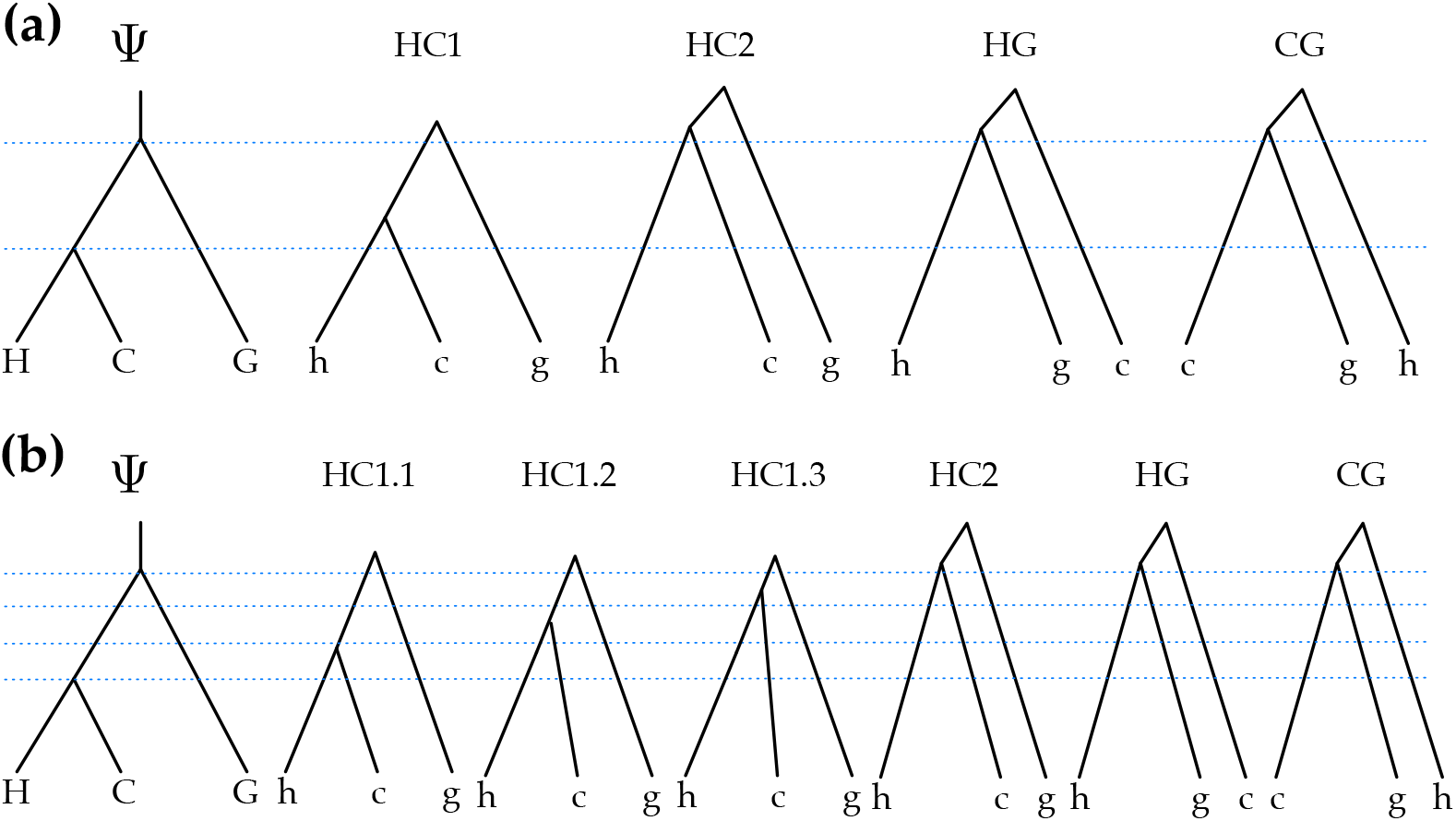
The coalHMM model. (a) The four states of a standard coalHMM that corresponds to the species tree Ψ. (b) The six states of a refined coalHMM that corresponds to the species tree Ψ when its internal branch is broken into three sub-branches.

Finally, once the coalHMM is built, **Forward** runs the Forward algorithm (Durbin et al., 1998) to compute 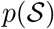 as an approximation of the likelihood 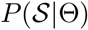.

### Bayesian Formulation and Variational Inference

As noted above, the data in our case is a sequence alignment 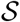 on the set of taxa of the species tree. We are interested in the posterior 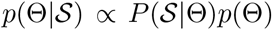, where we assume a prior distribution on Θ. Exact computation of the posterior is intractable, so we use variational inference to find an approximate distribution to 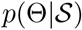. In variational inference, we posit a simple family of distributions over Θ and try to find the member of the family closest in terms of the Kullback-Leibler (KL) divergence to the true posterior 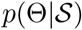 (Bishop, 2006). Denote the variational distribution we posit on Θ as *Q*(Θ|λ), governed by a set of free parameters λ, our goal is to approximate 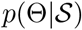 by optimizing λ to make *Q*(Θ|λ) as close in KL divergence to 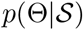 as possible. In variational inference, we optimize the Evidence Lower BOund (ELBO), given by

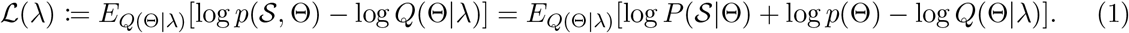

Maximizing 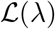 amounts to minimizing the KL divergence from *Q* to *p*. To maintain the general nature of VICAR and minimize the burden on users, we use Black Box Variational Inference (BBVI) (Ranganath, Gerrish, and Blei, 2014). BBVI is a stochastic optimization algorithm using noisy estimates of the gradient to maximize the ELBO, without the need for model-specific derivations (hence the “black box”). The gradient of the ELBO (Eq. (1)) with respect to λ can be written (Ranganath, Gerrish, and Blei, 2014) as

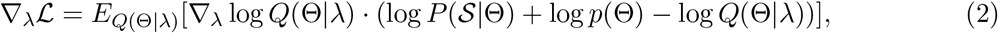

and its noisy unbiased Monte Carlo estimate is

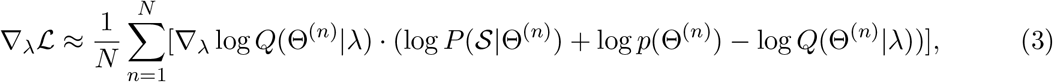

where Θ^(*n*)^ ~ *Q*(Θ|λ) is the *n*-th of *N* samples from the current variational distribution. All parts of the equations are known: ∇_λ_ log *Q*(Θ^(*n*)^|λ) is the score function (Cox and Hinkley, 1979) of the current variational distribution; an approximation of 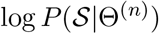 is computed by Algorithm 1 above; log *p*(Θ^(*n*)^) is the prior; and log *Q*(Θ^(*n*)^|λ) is computation about the variational distribution itself. Using Eq. (3) we can compute noisy gradients of 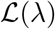 from samples of the variational posterior, and therefore we are able to do stochastic gradient ascent in the space of 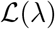 to optimize λ.

#### Factorized Approximation

As the variational family of *Q*(Θ), we assume the variational distribution to be a factorized Gaussian. Each parameter in Θ has a univariate Gaussian with a mean and a standard deviation. That is, each population size and node height of the species tree that we are interested in is independent and has a Gaussian variational posterior with its own mean and standard deviation. We have

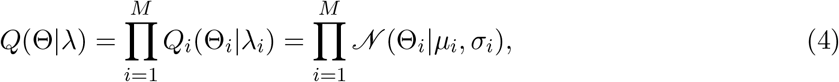

where *M* = |Θ| is the number of continuous parameters (divergence times and population sizes) associated with Ψ.

The per-component gradient of the ELBO with respect to each component of λ then becomes

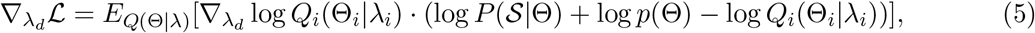

where λ_*d*_ belongs to the *i*-th factor of the factorized variational distribution. For factorized Gaussian, each factor has two λ components, mean and standard deviation. Taking all together, Algorithm 2 gives the general framework of VICAR. We note that under the BBVI framework, other variational families can serve as a drop-in replacement of the factorized Gaussian. Our method is easily generalizable to many other forms of variational distributions. In future implementations, we plan to support additional variational distributions so that users can choose a more flexible variational family and hence better approximation posteriors.

#### Variance Reduction and Adaptive Learning Rate

While Algorithm 2 gives the basic framework of the method, a few more challenges remain to be addressed to make it useful. In particular, the variance of the Monte Carlo estimator of the gradient given in Eq. (3) can be too large to be useful. To reduce the variance of the sampled estimator, we use control variates (Ross, 1997; Ranganath, Gerrish, and Blei, 2014). Control variate estimators are a family of functions with equivalent expectation but smaller variance than the function being approximated by Monte Carlo. Details of the application of control variates to BBVI can be found in (Ranganath, Gerrish, and Blei, 2014).

**Algorithm 2:**
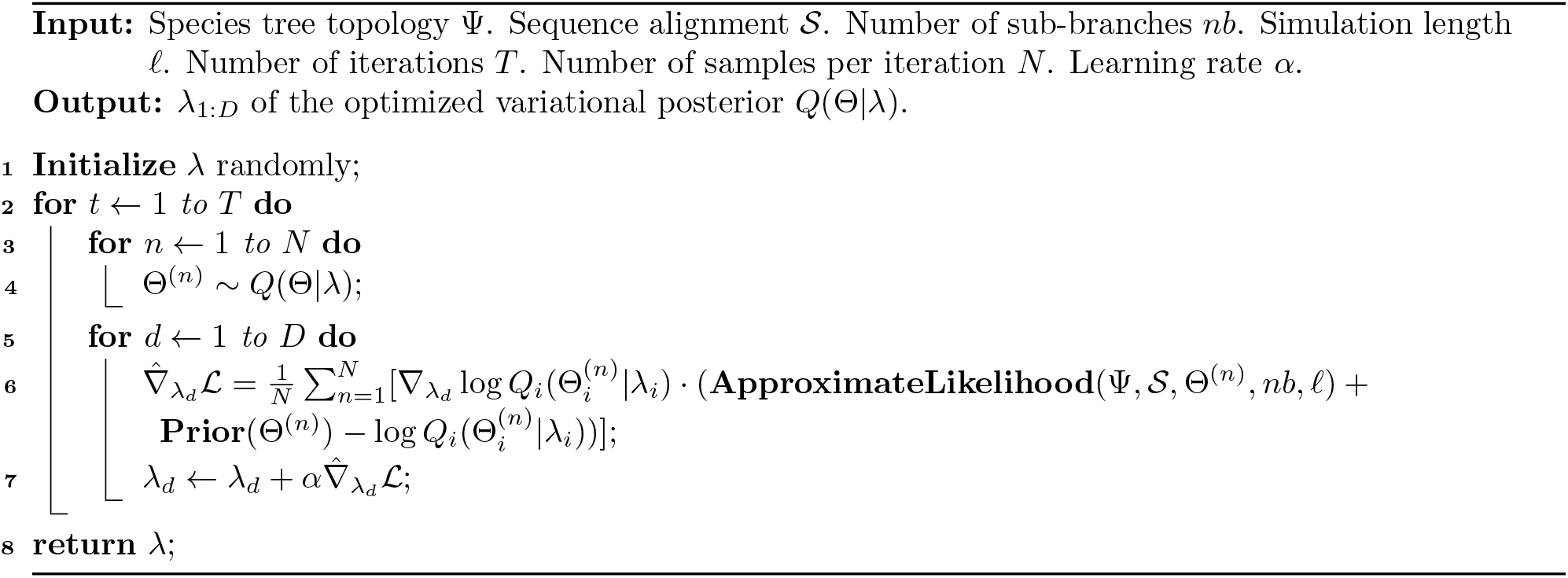
VICAR.

Another crucial challenge is setting the learning rate schedule. A large learning rate might overshoot the optimum, but a small learning rate might never converge. Moreover, the variational distribution in our problem has different scales (the scale of the population sizes and the node heights are different), so we would like our learning rate to be able to handle the smallest scale while not too small for the largest scale. As a result, we implement the AdaGrad (Duchi, Hazan, and Singer, 2011) optimizer to adaptively set the learning rate. AdaGrad adapts each learning rate by scaling it inversely proportional to the square root of the sum of all the squared past values of the gradient, resulting in greater progress in the more smoothly sloped direction of the parameter space and smaller progress otherwise. AdaGrad is a per-parameter updater, meaning it has a different adaptive learning rate for each parameter, addressing the multi-scale problem of our distribution. Other off-the-shelf optimizers like RMSProp (Tieleman and Hinton, 2012) and Adam (Kingma and Ba, 2014) can also be easily implemented within our framework.

### A Divide-and-conquer Approach to Larger Numbers of Genomes

While VICAR is general enough so as to handle an arbitrary number of taxa, its running time could increase super-exponentially in the number of taxa. It is important to note, though, that, in practice, this number could grow much slower, depending on the evolutionary parameters, which control the number of distinct states generated by the simulation underlying **ApproximateLikelihood**.

We propose a divide-and-conquer approach to ameliorate this problem. The method first divides the set of taxa into overlapping three-taxon subsets whose subtree parameters cover all the parameters of the full species tree, and then infers the parameters of each subtree using Algorithm 2.

For the divide step, in order to cover all the continuous parameters of the full species tree Ψ, we only need to cover all the internal edges. Using three leaves to cover an internal branch *e* = (*u, v*) in Φ, we need one leaf from the left child clade of *v*, one leaf from the right child clade of *v*, and one leaf from the right child clade of *u*. The process is illustrated in Fig. 15. The full species tree on taxa A, B, C, and D has two internal branches, each of which is covered by one of the two subtrees shown in the figure. There are a total of six continuous parameters for the full tree: node heights *T*_1_, *T*_2_, and *T*_3_, population sizes *N*_12_, *N*_23_, and root population size *N*_1_. Analysis of data from taxa *A*, *B*, and *C* allows for inferring the parameters *T*_2_, *T*_3_ and *N*_23_. Analysis of data from taxa *A, C*, and *D* allows for inferring the parameters *T*_1_, *T*_2_, *N*_12_, and *N*_1_.

**Figure 15:**
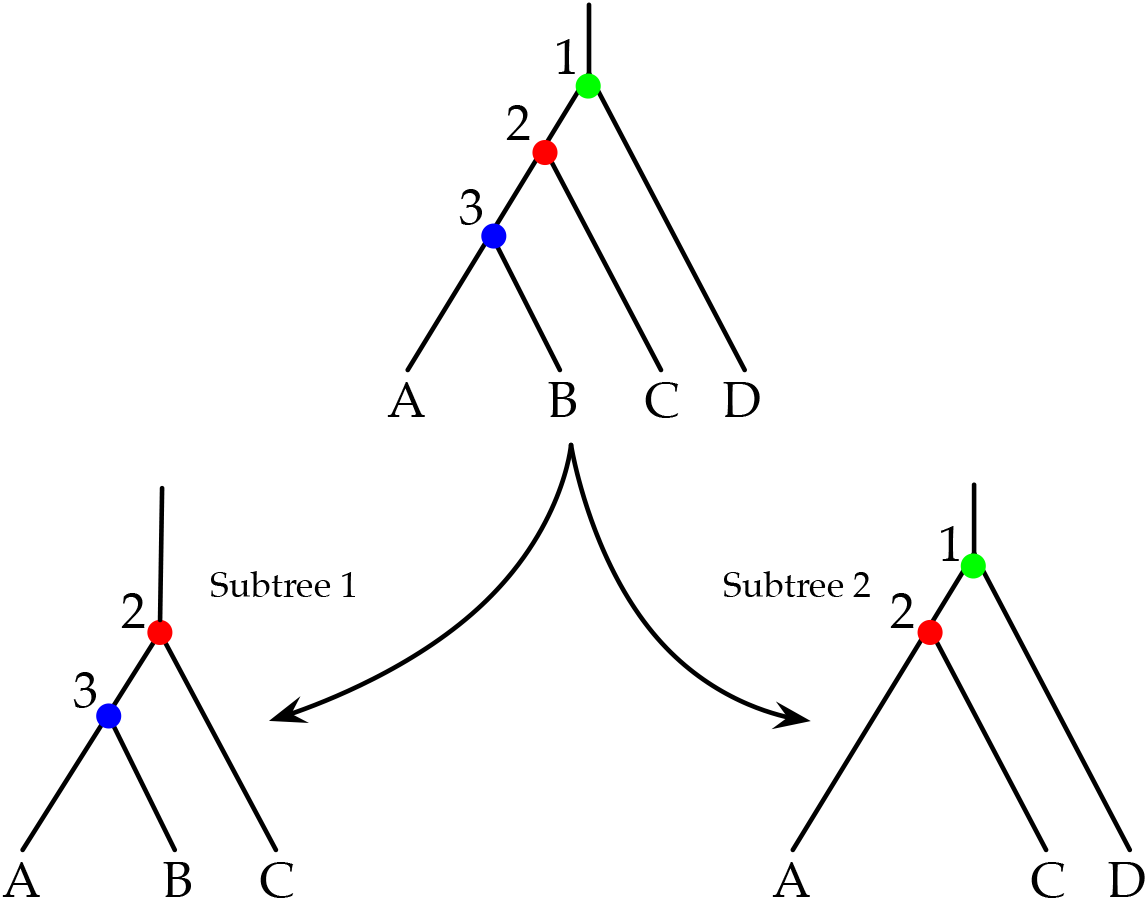
Divide-and-conquer inference on a four-taxon data set. The set of taxa {*A, B,C, D*} is divided into two sets {*A, B, C*} and {*A, C, D*} along with their respective species trees. Each of the two data sets are analyzed and the results are merged.

While in this work we consider each subset independently when running Algorithm 2, a future direction involves implementing an algorithm that is aware of the parameter overlap. For example, when the two data sets are analyzed, the algorithm is made aware of the fact that *T*_2_ is the same parameter in both data sets. This should scale linearly with the number of taxa, as we would need to infer the parameters for one triplet per internal branch.

## Software Availability

VICAR has been implemented in Java and is freely available as part of the software package PhyloNet (Than, Ruths, and Nakhleh, 2008; Wen et al., 2018).

## Acknowledgments

L.N. was supported by National Science Foundation Grants DBI-2030604, CCF-1514177, CCF-1800723, and DMS-1547433.

